# GBP2 engages Galectin-9 for immunity against *Toxoplasma gondii*

**DOI:** 10.1101/2021.12.28.474342

**Authors:** Elisabeth Kravets, Gereon Poschmann, Sebastian Hänsch, Veronica Raba, Stefanie Weidtkamp-Peters, Daniel Degrandi, Kai Stühler, Klaus Pfeffer

## Abstract

Guanylate binding proteins (GBPs) are large interferon-inducible GTPases, executing essential host defense activities against *Toxoplasma gondii,* an invasive intracellular apicomplexan protozoan parasite of global importance. *T. gondii* establishes a parasitophorous vacuole (PV) which shields the parasite from the host’s intracellular defense mechanisms. Murine GBPs (mGBPs) recognize *T. gondii* PVs and assemble into supramolecular mGBP homo- and heterocomplexes that are required for the disruption of the membrane of PVs eventually resulting in the cell-autonomous immune control of vacuole-resident pathogens. We have previously shown that mGBP2 plays an important role in *T. gondii* immune control. Here, to unravel mGBP2 functions, we report Galectin-9 (Gal9) as a critical mGBP2 interaction partner engaged for immunity to *T. gondii*. Interestingly, Gal9 also accumulates and colocalizes with mGBP2 at the *T. gondii* PV. Furthermore, we could prove the requirement of Gal9 for growth control of *T. gondii* by CRISPR/Cas9 mediated gene editing. These discoveries clearly indicate that Gal9 is a crucial factor for the mGBP2 coordinated cell autonomous host defense mechanism against *T. gondii*.

## Introduction

After innate immune sensing of pathogens via pattern recognition receptors, production of interferons (IFN) is a key prerequisite for the activation of cell-autonomous immunity, which is essential for the immediate restriction of microbial replication (1, 2). IFN-γ signaling induces the expression of hundreds of interferon-stimulated genes (ISGs) (3–5), which essentially contribute to innate and adaptive immune responses in immune and nonimmune cells (6, 7). Very prominent among IFN-γ ISGs are the members of several families of interferon-inducible GTPases such as the 65- to 73-kDa dynamin-related guanylate-binding proteins (GBPs) (8, 9) and the 45-kDa immunity-related GTPases (IRGs) (10, 11). Human (hGBPs) and murine GBPs (mGBPs) families consist of seven (hGBP1 to hGBP7) and eleven (mGBP1 to mGBP11) members, respectively, which are strongly upregulated in response to IFN-γ (12–14). GBPs characteristically exhibit various cellular functions encompassing antiparasitic, antiviral, and antibacterial activity, as well as inhibitory effects on cell proliferation and tumor suppression (15–26).

GBPs display a characteristic multidomain architecture including a large catalytic GTPase (G) domain and a C-terminal, helical domain (8, 27–33). Various mGBPs such as mGBP1, mGBP2, and mGBP7 are especially important effector molecules for the resistance against *Toxoplasma gondii*, an intracellular apicomplexan protozoan parasite of global relevance (16, 25, 34–39). After the active invasion of host cells, *T. gondii* forms a nonfusogenic cytoplasmic membranous structure called the parasitophorous vacuole (PV) in which the parasite efficiently replicates (37, 40, 41). mGBPs are intricately involved in IFN-γ induced growth restriction and killing of the *T. gondii* in murine macrophages, fibroblasts and astrocytes (16, 20, 25, 36, 38, 42). In order to execute their antimicrobial function, mGBPs can specifically associate with pathogen containing compartments of intracellular microbes which encompass pathogen-containing vacuoles (PCVs) as well as viral replication compartments (20, 25, 36, 43–45), resulting in the innate immune detection and cell-autonomous clearing or of PCV-resident pathogens (22, 25, 38, 46). After IFN-γ induction, mGBP2 molecules assemble in vesicle like structures (VLS) in the cytoplasm and, after *T. gondii* entry into the host cell, mGBP2 attacks the PV membrane (PVM) in orchestrated, supramolecular complexes, ultimately promoting the destruction of PVMs (36, 38). This stage is followed by recruitment of mGBP2 to the plasma membrane of the intracellular parasite, which in turn likely leads to its destruction (38). Multiparameter fluorescence image spectroscopy (MFIS) revealed that mGBP2, on a molecular level, can form homo- or heteromultimers with mGBP1, mGBP3 and, to a lesser extent, with mGBP5 within VLS and at the PVM (38). For PV membrane interactions, isoprenylation / farnesylation and/or hydrophobic moieties in the C-terminus of the GBPs are essential (25, 31, 32, 38, 47). Eventually, especially mGBP2 and mGBP7, can be found localized at the parasite cell membrane and inside the targeted parasite (25, 38) probably also interfering with the integrity of *T. gondii* membranes. Importantly, inactivation of mGBP1, mGBP2, or mGBP7 in mice has been shown to result in a vastly impaired resistance to infections with type II *T. gondii* (25, 36).

Galectins are a family of lectins characterized by a carbohydrate recognition domain (CRD) with β-galactoside-binding ability (48). Galectins are nucleocytoplasmic proteins, can be exported from cells through a poorly defined unconventional secretion pathway (49) and are able to directly bind to bacteria thereby mediating antibacterial effects and modifying inflammatory signaling events (50). Presently, 15 galectin members have been identified in mammals (51). Immunomodulatory effects of galectins include recruitment of immune cells to the site of infection, promotion of neutrophil function, and stimulation of the bactericidal activity of infected neutrophils and macrophages (48). Previously it has been shown that Gal3 can bind *T. gondii* glycophosphatidylinositols and might be a co-receptor presenting GPIs to TLRs on macrophages (52). Cytoplasmic galectins fulfill functions in several cellular programs including cell growth and apoptosis (53). Through binding to multivalent carbohydrates, galectins can multimerize and thereby cluster these surface glycoconjugates resulting in changes in cell migration, receptor endocytosis and various other cellular events (54, 55). Glycans are largely restricted to the luminal face but absent from the cytosolic face of intracellular vesicles. If vesicular damage occurs, luminal glycans are exposed to the cytosolic milieu, where they can be recognized by galectins (56, 57) . Furthermore, vacuolar instability of bacterial PCVs is recognized by host β-galactoside-binding proteins of the galectin family (50, 58). Intracellular galectins can serve as danger receptors and promote autophagy of the invading pathogen (59).

The aim of this study was to explore the molecular interaction(s) that enable mGBP2 to control *T. gondii* replication. By employing immunoprecipitation (IP) followed by mass spectrometry (MS) analysis, we have identified galectin-9 (Gal9) as a novel interaction partner of mGBP2. Furthermore, we demonstrate that the inactivation of Gal9 impairs the cell autonomous IFN-γ induced immune control of *T. gondii* replication.

## Results

### Identification of Gal9 as mGBP2 interaction partner by IP and MS

Previously, we could show that mGBP2 interacts with itself and other mGBP family members forming homo- and heteromultimers, respectively, in VLS and at the *T. gondii* PVM (38). For a profound understanding of the functions of mGBP2 in *T. gondii* infection, we searched for other interaction partners of mGBP2. To this end, mGBP2^-/-^ murine embryonic fibroblasts (MEFs)(36) expressing HA-tagged mGBP2 were constructed, stimulated with IFN-γ, and either infected with the type II strain *T. gondii* ME49 or left uninfected. An HA-tag specific IP was performed (Fig.S1A) followed by MS based quantitative protein analysis (Fig.1). mGBP2^-/-^ MEFs transduced with an empty expression vector served as negative controls.

**FIGURE 1.**
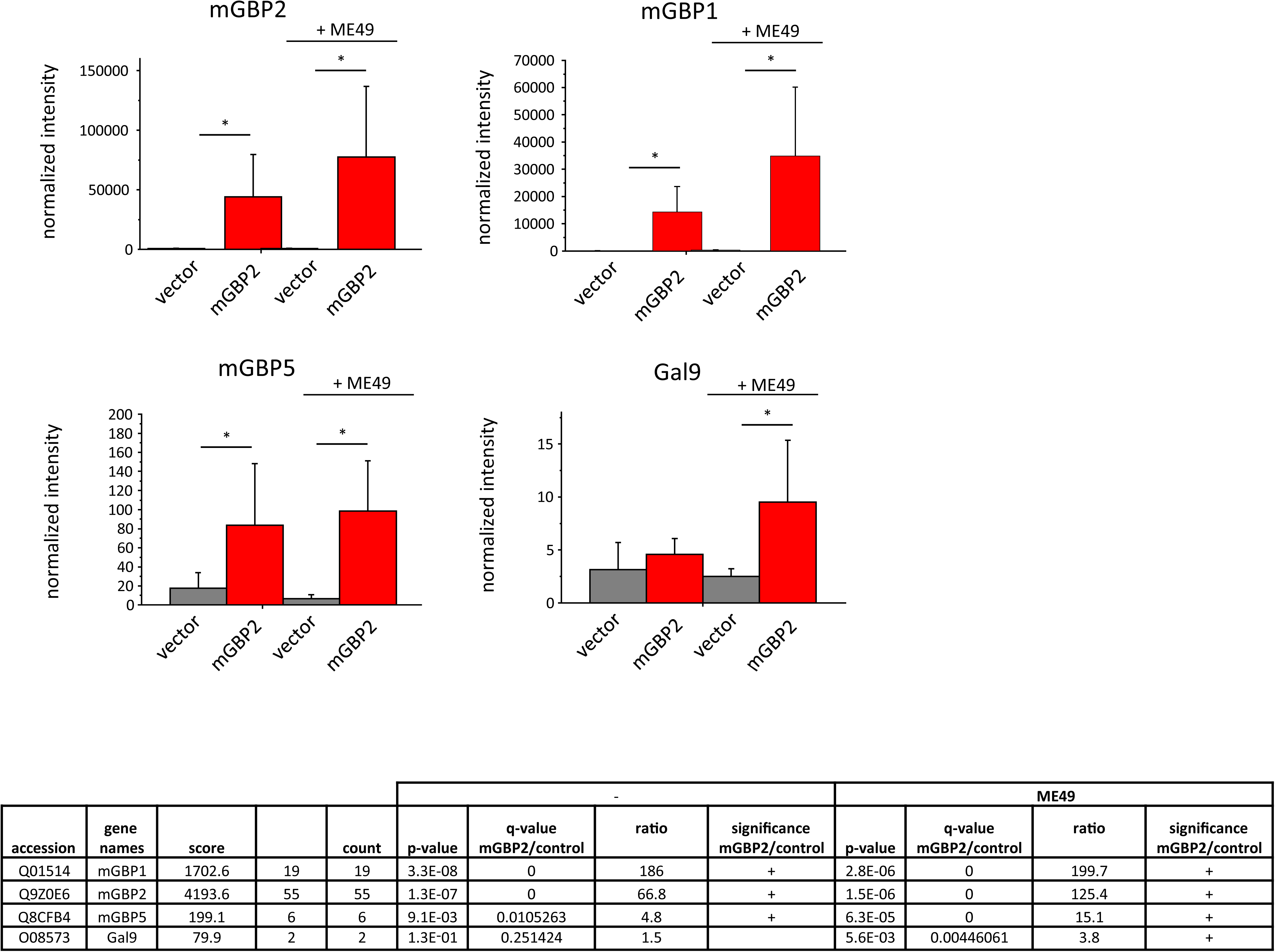
Identification of mGBP2 interacting proteins using co-immunoprecipitation in combination with label- free quantitative mass spectrometry. An HA-mGBP2 fusion protein was expressed in mGBP2^-/-^ MEFs. mGBP2^-/-^ MEFs transduced with the empty expression vector were used as controls. MEFs were stimulated with IFN-γ (16h) and left uninfected or were infected with *T. gondii* ME49 for 2h. IPs were performed with an anti-HA antibody from cell lysates. Immunoprecipitated proteins were analyzed by label-free quantitative mass spectrometry. Five to six replicates were analyzed per condition and putative mGBP2 interacting proteins were identified from uninfected and *T. gondii* ME49 infected cells. Mean values of normalized intensities are shown in the bar plots ± standard deviations. In the table, details from the protein identification are shown: the confidence scores from the MASCOT search engine (75, 76), the number of identified unique peptides per protein as well as p-values from Student’s t-tests. Furthermore, the enrichment ratios (intensity ratio between mGBP2 expressing cells and vector control) are given (a correction for multiple testing has been performed using the significance analysis of microarray approach, 5% false discovery rate, S0 0.8, a minimum of 4 valid values per group).

Statistical analysis revealed that 148 proteins were co-purified with HA-mGBP2 from the uninfected and 294 proteins from the *T. gondii* infected cells. Besides the bait protein mGBP2, further members of the mGBP family were enriched as prey proteins. Among them, a clear co-purification of mGBP1 and mGBP5 which previously were independently identified by MFIS as mGBP2 interacting members, could be observed in uninfected and infected HA-mGBP2 expressing MEFs (mGBP1 enrichment factor > 185 in both experimental conditions, mGBP5 enrichment factor 4.8 from uninfected and 15.1 from infected cells) (Fig.1), verifying the reliability of the co-IP and MS analysis.

From the list of potential mGBP2 interactors, Gal9 was analyzed further since Gal9 was found significantly enriched in IP samples from type II *T. gondii* ME49 infected cells (Fig.1) and interestingly a role for several other galectin family members had already been proposed in microbial infections (51).

To validate and characterize the interactions of Gal9 with mGBP2 in detail, mCherry-fused proteins of Gal9 were co-expressed with full-length mGBP2 or with mGBP2 truncation mutants containing the GTP-binding domain and the middle domain (GM), the middle and the C-terminal effector domain (ME) or the C-terminal effector domain only (E) (31, 36). These cell lines were infected with *T. gondii* followed by co-IP. Using this approach, we could prove the interaction of mGBP2 with Gal9 and could determine a most prominent function of the E-domain of mGBP2 for the interaction with Gal9 (Fig.2 and Fig. S1B).

**FIGURE 2.**
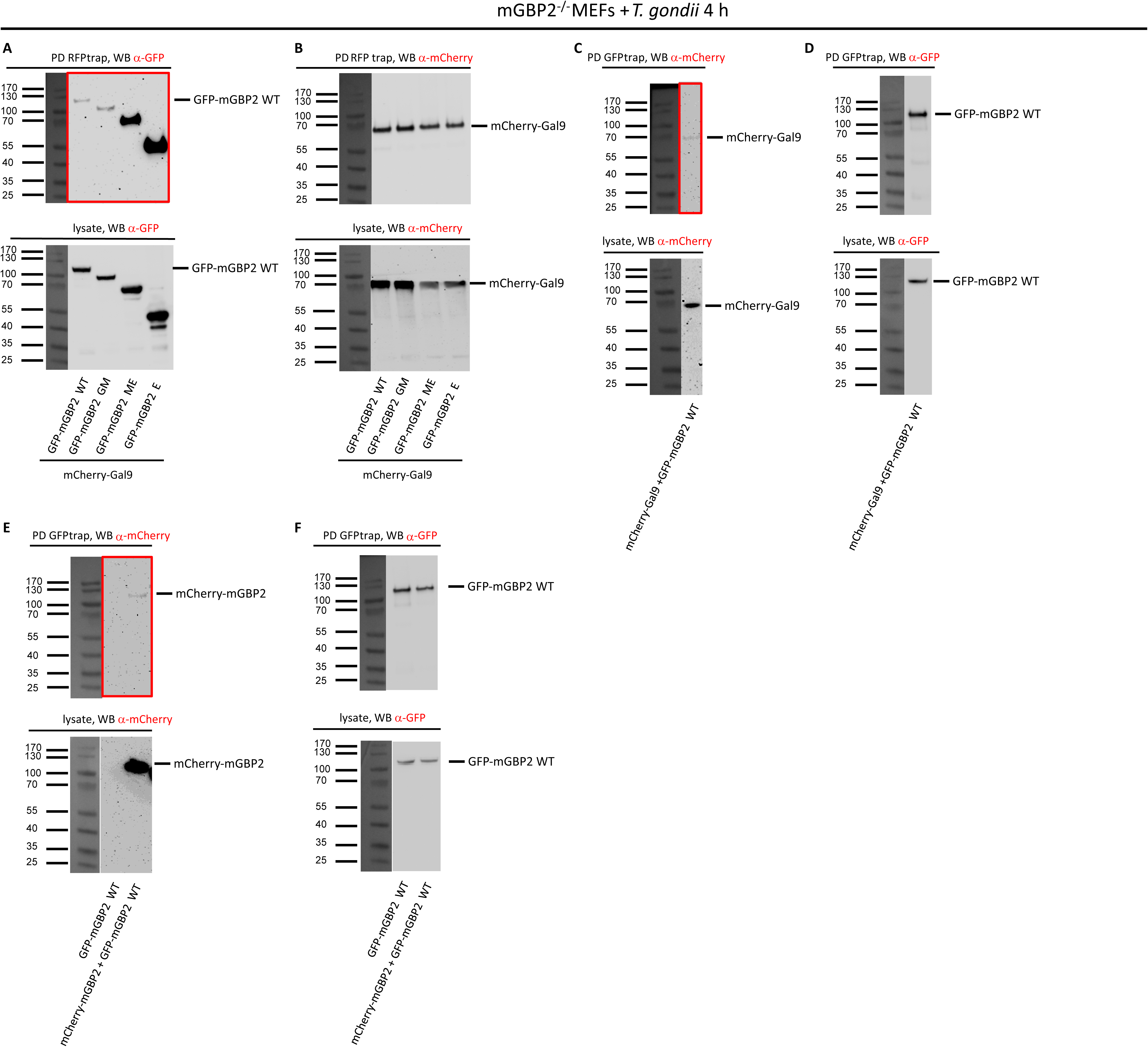
Verification of Gal9 as interaction partner of mGBP2 by pulldown analysis. mGBP2^-/-^ MEFs were reconstituted with GFP-mGBP2 WT or one of the indicated mGBP2 truncation mutants as well as one of the N-terminal mCherry fusion proteins mCherry-mGBP2 or mCherry-Gal9. 10^6^ Cells were stimulated with IFN-γ for 16 h and infected with *T. gondii* ME49 for 4h, MOI 50. Cells were lysed and lysate supernatants were incubated o/n with either RFP-Trap^®^ beads (A, B) or with GFP-Trap^®^ beads (C-F) at 4°C. Pull-down (PD) samples and appropriate cell lysate supernatants were subjected to Western Blotting (WB). Blots were stained with α-GFP (A, D, F) or α-mCherry (B, C, E) antibodies. GTP-binding domain (GM), the middle domain and the C-terminal effector domain (ME) or the C-terminal effector domain only (E) (36).

### Subcellular localization of Gal9 and recruitment to the T. gondii PV

In a next step, the colocalization of mGBP2 and Gal9 at the PVM of *T. gondii* was determined by confocal (Fig.S2) and Stimulated Emission Depletion (STED) microscopy (Fig.3). Doubly transduced mGBP2^-/-^ MEFs coexpressing GFP- mGBP2 with mCherry-Gal9 were prestimulated with IFN-γ and infected with *T. gondii* ME49 (Fig.3, Fig.S2). GFP- mGBP2 and mCherry-mGBP2 coexpressing MEFs served as a positive control in STED microscopy colocalization experiments (Fig.S3). A distinctive colocalization or close vicinity between mGBP2 and Gal9 could be observed at the PVM already 2 h after infection. Interestingly, mCherry-Gal9 partly colocalized in GFP-mGBP2 containing VLS in uninfected MEFs (Fig.S4). Furthermore, 5-6 h after infection Gal9 and mGBP2 could be found to be located together within the intermembranous space of the PV and eventually within the cytosol of the parasite (Fig.S2). These results indicate that Gal9 is interacting and acting together with mGBP2 and suggest that Gal9 is intricately involved into the cell autonomous immunity mediated by mGBP2.

**FIGURE 3.**
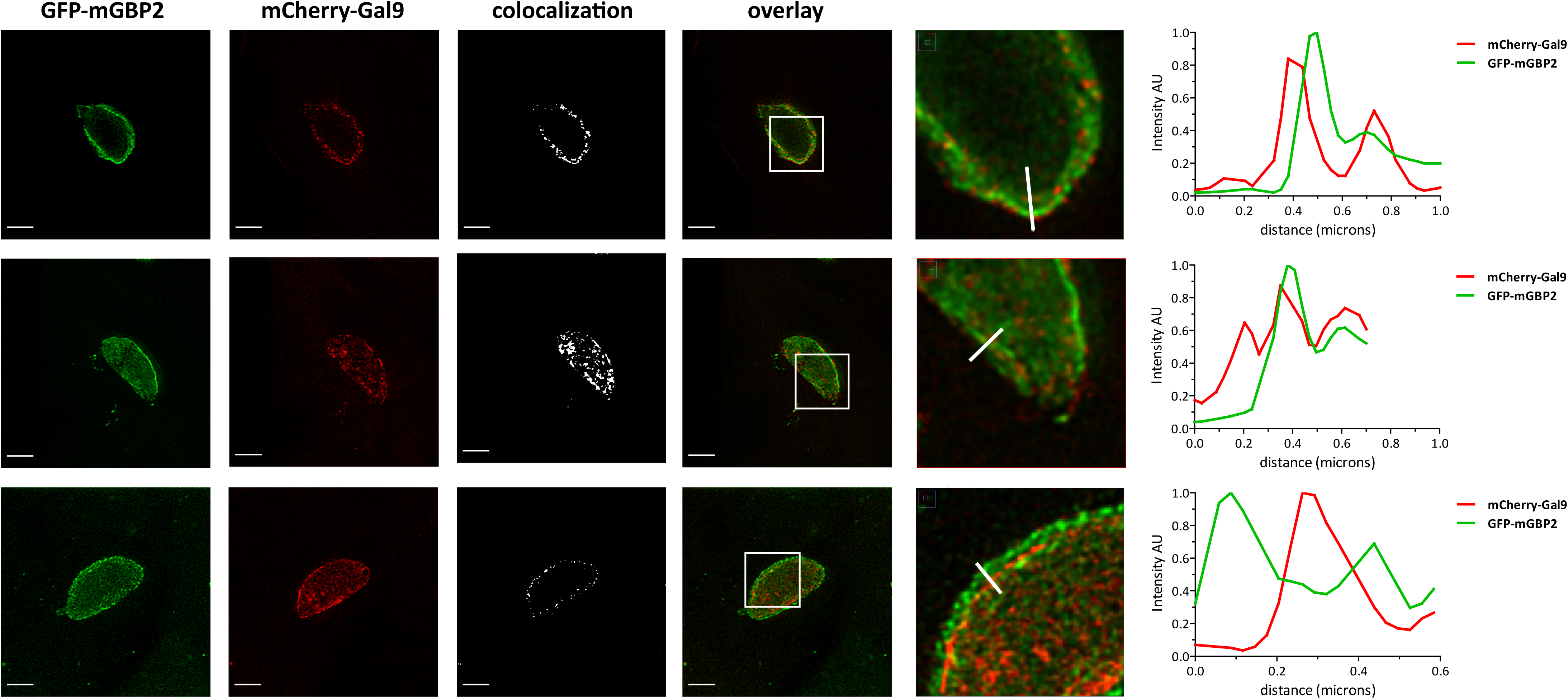
Intracellular colocalization of mGBP2 interaction partner Gal9 at the PVM of *T. gondii*. Recruitment and colocalization of mGBP2 and Gal9 was analyzed after transduction of a GFP-mGBP2 fusion construct in mGBP2^- /-^ MEFs and additional transduction of mCherry-Gal9. MEFs were seeded and incubated on glass slides, stimulated with IFN-γ for 16 h and subsequently infected with *T. gondii* ME49 for 2 h. After fixation, infected cells were treated with an α-RFP VHH nanobody conjugated to eGFPBoosterAtto647N and with an α-GFP VHH nanobody conjugated to eGFPBoosterAtto488 for enhancement of the immunofluorescence of mCherry and GFP, respectively. Glass slides were analyzed by STED microscopy. Bars 2 µm. The graphs in the right panel depict a fluorescence intensity analysis of STED images on the far right with the ImageJ software (Fiji) for Atto488 and Atto647 fluorescence signals along the cross sections of PVMs as indicated. The colocalization thresholds were set to 7500-max for the Cherry and 5000-max for GFP in mCherry-Gal9 and GFP-mGBP2 expressing cells.

### T. gondii replication control by Gal9

Next, the contribution of Gal9 to cell autonomous immunity against *T. gondii* was investigated. For this purpose, the *lgal9* gene was inactivated applying CRISPR/Cas9 gene editing. The inactivation of Gal9 in different clonal NIH 3T3 fibroblast cell lines was confirmed by sequencing of the targeted gene loci and, additionally, by Surveyor nuclease assay and WB for Gal9 (Fig.S5). Gal9- deficient fibroblast cell lines were stimulated with IFN-γ and infected with *T. gondii* (ME49) and the replication of the parasite was monitored. First, it was determined whether the inactivation of Gal9 leads to increased or decreased infection rates in the targeted cell lines compared to wildtype cells. However, the number of *T. gondii* parasites detected in the gene edited cell lines compared to wildtype fibroblasts did not differ 2 h post infection (Fig.S6). Interestingly, a significant increase in *T. gondii* replication could be observed in fibroblasts harboring an inactivation of Gal9 as determined by qPCR and by quantification of the number of *T. gondii* rosettes and single parasites 24 h after infection (Fig.4, Fig.S7. To assess whether the differences in *T. gondii* control could be attributed to altered cell death, we performed LDH release assays of WT and Gal9 deficient cells including cells reconstituted with the appropriate construct. We could not observe any difference in cell death in Gal9-deficient cells 24h after infection (Fig. S8).

**FIGURE 4.**
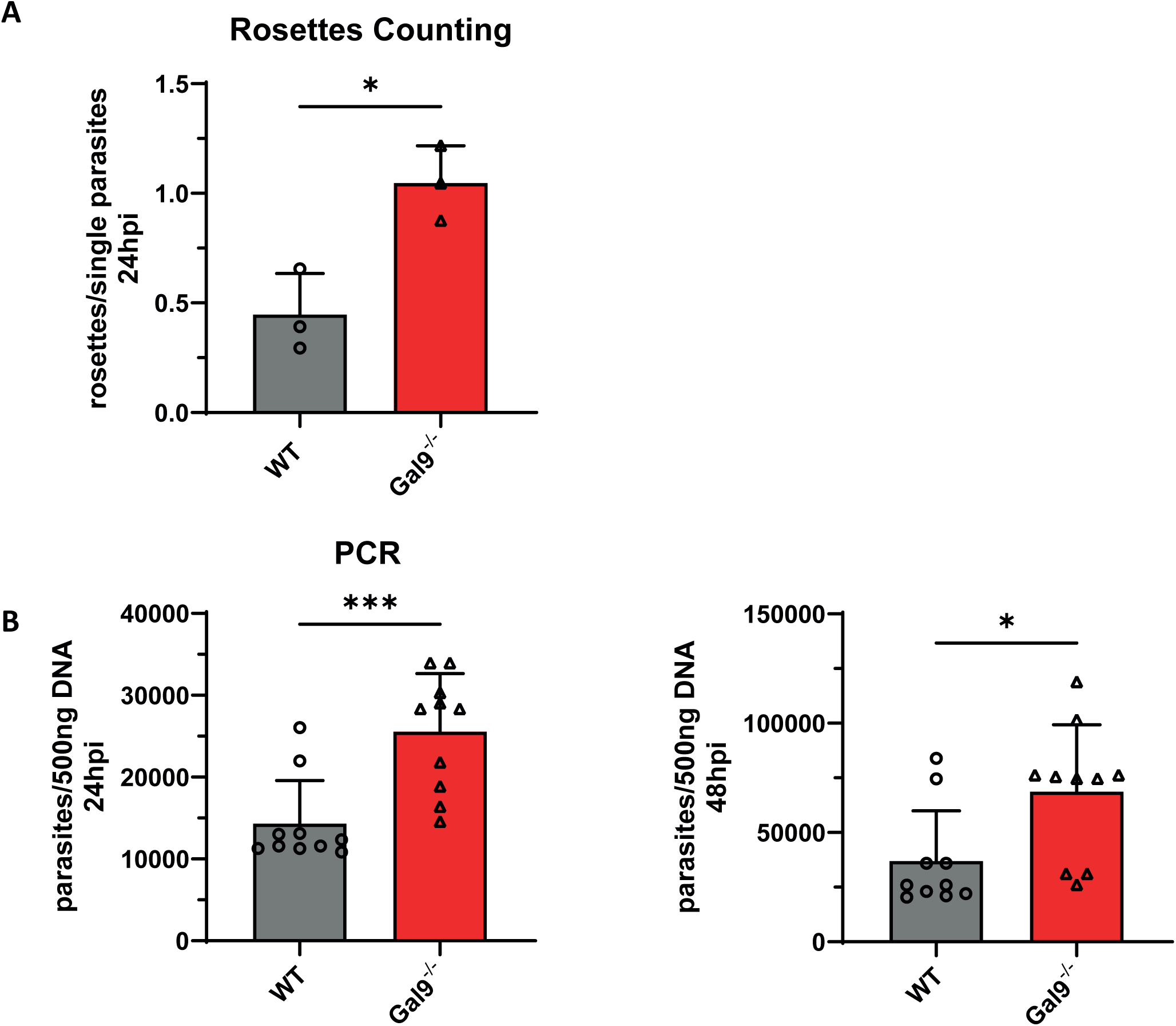
Gal9 is required for the intracellular control of replication of *T. gondii*. (A) WT and NIH 3T3 cell line clones with verified CRISPR/Cas9 mediated inactivation of Gal9 (see main text and Figs.S5/S6) were stimulated with IFN-γ for 16 h and subsequently infected with *T. gondii* ME49 MOI10 for 24 h. After fixation, *T. gondii* were stained with an α-SAGI antibody and the cell nuclei were labeled with DAPI. Glass slides were analyzed by confocal microscopy. The amounts of rosettes (replicative units) and single parasites inside the PV were determined in WT, Gal9- deficient NIH 3T3 fibroblasts 24 h after infection. In each cell line more than 100 PVs were counted in 3 independent experiments. Mean values ± SD are shown. (B) WT and Gal9 deficient NIH3T3 cells were stimulated with IFN-γ for 16 h and subsequently infected with *T. gondii* ME49 MOI10 for 24 and 48h. Genomic DNA from these samples was extracted and a *T. gondii* specific qPCR was performed. Parasite equivalents per 500 ng DNA are depicted for at least 10 individual samples from three independent experiments. Statistical analysis was performed using the Student’s t-test. ***: p <= 0.001; **: p <= 0.01; *: p <= 0.05.

These results clearly indicate an important role for Gal9 in controlling the intracellular replication of the parasite and suggest that mGBP2 and Gal9 function as supramolecular complexes in the defense against intracellular pathogens.

## Discussion

mGBP2 provides cell-autonomous immunity to the intracellular PV resident parasite *T. gondii* (31, 36, 38, 44, 60). Although it has been shown that mGBPs translocate to PVs, form supramolecular complexes, attack the membrane of the parasite itself, and, consecutively, restrict replication of the intracellular pathogen, the molecular mechanisms by which mGBPs control the parasite inside the host cell remain largely elusive. Here we identify a novel interaction partner of mGBP2, Gal9, which plays a critical role with respect to the recruitment and in the antiparasitic function of mGBP2 in the context of cell-autonomous defense against *T. gondii*.

Previously, it was shown that Gal8 colocalizes with mGBP2 positive *Salmonella spp*. in unprimed bone marrow derived macrophages (BMDMs) (61). It was proposed that Gal8 monitors endosomal and lysosomal integrity and detects bacterial invasion of the cytosol by *Listeria* spp. or *Shigella spp.*, serving as a versatile receptor for vesicle-damaging pathogens (61). Interestingly, it was also reported that Gal3, as a member of the Galectin family, and mGBP2 form protein complexes, which associate with vacuoles harboring secretion system-competent *Legionella pneumophila* and *Yersinia pseudotuberculosis* bacteria in response to vacuolar damage (62). In this study we show that, either directly or indirectly, Gal9 interacts and colocalizes with mGBP2 in IFN-γ stimulated MEFs and NIH 3T3 cells in VLS. Considering the methodology used in this study, the possibility exists that Gal9 does not physically interact with mGBP2, but that other proteins/molecules in a complex may be involved in bridging the interaction. However, this would not affect the overall significance of the study. Based on its structural organization, Gal9 belongs to the tandem-repeat type galectins containing two distinct but homologous CRDs (N-CRD and C-CRD) covalently connected by a linker peptide (48, 59). Recruitment of Gal9 to the *T. gondii* PV has not been described previously. Furthermore, Gal9 could not only be found at the PVM but also within the intermembranous space between the PVM and the *T. gondii* plasma membrane as well as in the cytosol of the parasites together with mGBP2 approx. 5-6 h post-infection. This indicates that Gal9 and mGBP2 might recruit to the PVM of *T. gondii* in preformed complexes. Mechanistically, a series of events involving mGBP/IRG mediated PVM indentation, PVM vesiculation/disruption, and parasite plasma membrane stripping, leading ultimately to degradation of the parasite, is proposed by us and others as hallmarks for intracellular *T. gondii* elimination (38, 63, 64). It can be suggested that pore forming or denudating activity of these IFN-γ induced GTPases allows Gal9 to transmigrate through the PVM and to recognize β-galactosides on the inner leaflets of the PVM and the parasite, labeling the parasite for further degradation processes. Parasite glycoconjugates play important roles in host cell invasion, and specific interactions between host galectins and parasite glycoconjugates are considered to be critical for pathogen recognition (55). Interestingly, glycosylphosphatidylinositols (GPIs) of *T. gondii* (65) have been shown to be ligands of human Gal3 whereby both glycan and lipid moieties participate in the association of *T. gondii* GPIs with Gal3 (52). In this context, Gal3, -8, and -9 bind to host glycans in the luminal side of lysed phagosomes or permeated pathogen vacuoles (66). Intracellular Gal3 was characterized as a sensor of vacuolar rupture that occurs when bacteria such as *Shigella flexneri* or *Salmonella enterica* serovar Typhimurium actively enter the host cell cytosol (58, 67). Gal3, -8 and -9 were also shown to associate with Salmonella-containing vacuoles (SCV) (61, 68, 69). Here we show that Gal9 is required for the control of the parasite *T. gondii*.

Functionally, the recruitment of mGBP2 to the PVM was reduced in Gal9- inactivated cells and vice versa; the recruitment of Gal9 to the PVM was reduced in mGBP2 deficient cells. In accordance, significantly fewer Salmonella were targeted by Gal8 in Gbp^chr3^ KO BMDMs than in wild-type BMDMs (70). Also, the delivery of mGBP1 and mGBP2 to Legionella-containing vacuoles (LCV) or Yersinia-containing vacuoles is substantially diminished in Gal3- and, to a lesser extent, Gal8-deficient IFN-γ–primed iBMDM (62). In Salmonella-infected cells, SCV rupture and Gal3 recruitment was followed by a rapid and massive recruitment of hGBP1 (67). Interestingly, a specific polybasic protein motif (PBM: KMRRRK) was identified for hGBP1 which is sufficient to drive significant targeting of hGBP1 to *S. flexneri* recognizing exposed glycans from disrupted, bacteria containing, endosomes. Deletion of the PBM-motive leads to substantially reduced colocalization between Gal3-marked vacuoles and hGBP1 (71). mGBP2 lacks such a C-terminal PBM. Nevertheless, the interaction with Gal9 appears to be mostly dependent on the C-terminal domain of mGBP2. Rab GDP dissociation inhibitor α (RabGDIα) acts as a suppressor of the mGBP2-Irga6 axis, hereby the lipid-binding activity of RabGDIα is critical for IFN-γ–induced mGBP2-dependent cell-autonomous immunity against ME49 *T. gondii*. RabGDIα deficiency is accompanied by increased recruitment of mGBP2 and Irga6 to the parasite in MEFs and macrophages and results in enhanced IFN-γ–mediated *T. gondii* clearance *in vitro* and *in vivo* (72). These studies suggest on the one hand GBPs as escorts for distinct anti-pathogenic factors en route to PVs, on the other hand that the recruitment of galectins and GBPs to the PVs might be mutually dependent as part of a coordinated host defense program. This is supported by the significantly increased *T. gondii* proliferation in IFN-γ stimulated Gal9 knock out 3T3 fibroblasts 22h post-infection indicating an important role of Gal9 in cell-autonomous *T. gondii* elimination. In contrast, IFN-γ–primed Gal3^−/−^ BMDMs restricted Yersinia and Legionella bacterial growth, similar to WT BMDMs (62).

Thus, it might be speculated that the recognition of the lysed PV by Gal9 initiates the uptake of the parasite into autophagosomes or other degradation pathways as it was described for *Salmonella*, where GBPs promote LC3 recruitment via Gal8 (70). Interestingly, *T. gondii* infected Gal3^−/−^ mice develop reduced inflammatory responses in many organs (73) and succumb to an intraperitoneal infection with *T. gondii* associated with a deficient influx of neutrophils and macrophages into the peritoneal cavity (74). The function of Gal9 might be the non-redundant to Gal3 and Gal8 in parasitic degradation processes which will be addressed in the future in a Gal9-gene deficient mouse line.

The findings of this study indicate that mGBP2 acts as a scaffold organizing the cell autonomous immunity in concert with Gal9 and thus extends the understanding of molecular host responses in the control of intracellular parasites.

## Ethics statement

Not applicable.

## Author Contribution statement

EK designed and performed most experiments; GP and KS performed and analyzed the mass spectrometry data; SH and SWP performed STED microscopy and analysis; VR performed LDH-release assays. DD and KP designed and supervised the study. EK, DD and KP wrote the paper. All authors contributed in writing, discussing and editing of the manuscript.

## Data availability statement

The mass spectrometry proteomics data have been deposited to the ProteomeXchange Consortium via the PRIDE (107) partner repository with the dataset identifier PXD027029.

## Acknowledgements

We thank Julia Mock, Nicole Küpper and Karin Buchholz for excellent technical assistance. We would like to acknowledge the Center for Advanced Imaging (CAi) at Heinrich Heine University for providing access to the Leica TCS SP8 gSTED 3X microscope system (DFG INST 208/665-1 FUGG).

## Experimental Procedures

### Expression Constructs

The WT ORF of mGBP2 (NCBI accession numbers: mGBP-2, NM_010260.1) was cloned either into the pWPXL plasmid (Trono lab) as N-terminal GFP-fusion constructs or into the pWPI Plasmid (Trono lab) in frame with a N-terminal HA-tag. For the cloning of mCherry constructs (36), the pWPXL plasmid was modified by exchanging of the gene for GFP by the gene for mCherry (38). mRNA was isolated from freshly prepared WT MEFs prestimulated with IFN-γ and infected with *T. gondii*. The ORFs of Galectin 9 (NM_010708.2) was amplified by PCR and then cloned into the modified pWPXL plasmid as N-terminal mCherry-fusion constructs (38). The lentiviral envelope vector pLP/VSVG (Invitrogen) and the packaging vector psPAX2 (Trono lab) were used for the lentiviral genetic transfer.

### Cell culture and transduction

NIH 3T3 fibroblasts (Public Health England; AccNo 93061524) and MEFs were cultured in Dulbecco’s modified Eagle’s medium (DMEM, Invitrogen/Gibco) supplemented with 10% (v/v) heat-inactivated low endotoxin fetal bovine serum (FBS, Cambrex), 100 U/ml penicillin, 100 μg/ml streptomycin, 2 mM L-glutamine (Biochrom) and 0.05 mM β- mercaptoethanol (Invitrogen/Gibco). Human foreskin fibroblasts (HS27, ATCC CRL-1634) were hold in culture in Iscove’s Modified Dulbecco’s Medium (IMDM, Invitrogen/Gibco) with the same supplementations. 293FT cells (Gibco, ThermoFisher; R700-07) were cultivated in DMEM supplemented with 10% FBS, 100 U/ml penicillin, and 100 μg/ml streptomycin. All recombinant lentiviruses were produced by transient transfection of 293FT cells according to jetPRIME^®^ transfection protocol (Polyplus, New York City) (38). NIH 3T3 cells and MEFs were seeded in 24 well plates (Corning Incorporated, Germany) and transduced with 600 µl of lentivirus with 25 µg Polybrene (Merck Millipore, Germany). After 5 h of incubation medium was changed. The transduction efficacy was analyzed by flowcytometry. Subsequently, only GFP or double GFP and mCherry positive cells were sorted and cultivated.

### In Vitro Passage of Toxoplasma gondii

Tachyzoites from *T. gondii* strain ME49 (ATCC 50611) were maintained by serial passage in confluent monolayers of HS27 cells. After infection of fibroblasts, parasites were harvested and passaged as described previously (16).

### Infection of murine MEFs with T. gondii

Cells were stimulated with 200 U/mL IFN-γ (R&D Systems, Minneapolis, MN) 16 h before infection. For immunofluorescence, NIH 3T3 fibroblasts and MEFs were cultured in 24-well plates (Falcon, BD Biosciences, Germany) on cover slips (ø 13 mm, VWR International, Germany) and inoculated with freshly harvested *T. gondii* at a parasites/cell ratio of 10:1. To remove extracellular parasites, cells were washed with PBS.

### Pull-Down and co-immunoprecipitation analysis of mGBP2

For co-immunoprecipitation experiments, mGBP2^-/-^ MEFs reconstituted with HA-mGBP2 were used. mGBP2^-/-^ MEFs transduced with the empty pWPI vector were used as control cells. 10^6^ Cells were then stimulated with IFN-γ for 16 h, infected with the *T. gondii* strain ME49 (MOI50) for 2 h or left uninfected, subsequently lysed and lysate supernatants were incubated o/n with monoclonal mouse anti-HA- antibody coupled to agarose beads (clone HA-7, Sigma-Aldich, USA) at 4°C. IP probes were subjected to silver staining and to Western Blotting. Blots were stained with the mGBP2- specific antiserum (Eurogentec), mouse α-GFP (clones 7.1 and 13.1, Roche, Switzerland) and rabbit αHA (polyclonal, Sigma-Aldrich, USA) antibodies. For pull-down experiments, lysate supernatants of mGBP2^-/-^ MEFs reconstituted either with WT GFP-mGBP2 or GFP-fused truncated mutants mutants or just transduced with a GFP expressing vector, as an appropriate control and coexpressing mCherry fused with Gal9 or just the mCherry protein, were incubated o/n with RFP-Trap^®^ or GFP-Trap^®^ beads (ChromoTek GmbH, Germany) at 4°C. IP probes were subjected to Western Blotting. Blots were stained with the mouse α-GFP or α-mCherry (Clontech, EU) antibodies.

### MS Analysis of interaction partners of mGBP2

A label-free mass spectrometry, based quantification approaches enabled by a highly reproducible and stable LC- MS/MS system was chosen for a quantitative analysis of the mGBP2 interactome. Six individual replicates of co- immunoprecipitation samples per group (HA-mGFP2, control-vector, HA-mGFP2 + ME49 and control-vector + ME49) were prepared from transfected IFNγ stimulated mGBP2^-/-^ MEFs (see above). Samples were prepared for mass spectrometric analysis essentially as described elsewhere (77). Briefly, proteins were stacked into an acrylamide gel (about 4 mm running distance), subjected to silver staining, de-stained, reduced and alkylated and digested with trypsin. Resulting peptides were extracted from the gel and approximately 500 ng peptides prepared in 0.1% trifluoroacetic acid for liquid chromatography and mass spectrometric analysis.

First, peptide samples were separated on an UltiMate 3000 RSLCnano chromatography system (Thermo Fisher Scientific). Peptides were initially trapped on a 2 cm long trap column (Acclaim PepMap100, 3 µm C18 particle size, 100 Å pore size, 75 µm inner diameter, Thermo Fisher Scientific) for 10 min at a flow rate of 6 µl/min with 0.1 % (v/v) TFA as mobile phase. Subsequently, they were separated on a 25 cm long analytical column (Acclaim PepMapRSLC, 2 µm C18 particle size, 100 Å pore size, 75 µm inner diameter, Thermo Fisher Scientific) at 60°C using a 2 h gradient from 4 to 40% solvent B (solvent A: 0.1% (v/v) formic acid in water, solvent B: 0.1% (v/v) formic acid, 84% (v/v) acetonitrile in water) at a flow rate of 300 nl / min.

Separated Peptides were injected with distal coated SilicaTip emitters (New Objective, Woburn, MA, USA) into an Orbitrap Elite mass spectrometer via a nanosource electrospray interface. The mass spectrometer was operated in positive, data dependent mode with a spray voltage of 1.4 kV and capillary temperature of 275°C. First, a full scan from m/z 350 to 1700 was recorded with a resolution of 60000 (at 400 m/z) in the obitrap analyser with a target for automatic gain control set to 1000000 and a maximal ion time of 200 ms. Then, up to 20 two- and threefold charged precursor ions with a minimal signal of 500 were isolated (isolation width 2) in the linar ion trap part of the instrument. Here, ions were collected for a maximum of 50 ms with an automatic gain control set to 30000, fragmented by collision induced dissociation with a normalized collision energy of 35 and analysed at a resolution of 5,400 (at 400 m/z). Already fragmented precursors were excluded from fragmentation for the next 45 seconds.

Quantitative analysis of MS1 based precursor ions was carried out with Progenesis QI for proteomics (version 2.0, Nonlinear Dynamics, Newcastle upon Tyne, UK). The software first detects and aligns precursor ion signals (features) then associates information from MS2 based precursor identification and finally computes protein intensities on basis of associated features. Automatic processing was used without manual alignment adjustment. Here, one run of the control-vector + ME49 could not automatically aligned and was not further considered in the analysis. Searches were triggered by Proteome Discoverer (version 1.4.1.14, Thermo Scientific) and the percolator node was used for identification validation at 1% false discovery rate. Spectra were searched with the Mascot search engine (version MASCOT 2.4.1, Matrix Science, London, UK) considering cysteine carbamidomethylation as fixed and protein N- terminal acetylation and methionine oxidation as variable modification as well as tryptic cleavage specificity with a maximum of two missed cleavage sites. Mass precision was set to 10 ppm for precursor and 0.4 Da for fragment spectra. Database searches were carried out using the mus musculus (UP000000589) and toxoplasma gondii (UP000002226) reference proteome data sets downloaded on 18^th^ January 2018 from UniProtKB including 52548 respectively 8404 entries. Only “high confident” peptide spectrum matches were used for further processing and peptide spectrum matches associated with proteins identified with at least two unique peptides. As we noticed a bias towards higher protein intensities in the mGBP2 overexpressing cells (arising from mGBP2 and associated proteins), no global normalization was performed but a normalization of protein intensities based on the sum of the Ig heavy chain proteins P18525, P18524. Calculated protein intensities were further processed within the Perseus (version 1.6.2.2, MPI for Biochemistry, Planegg, Germany) software environment. Here, Student’s t-tests were calculated on log2 transformed normalized intensities using the significance analysis of microarrays method for cutoff determination (S0 set to 0.8, 5% false discovery rate, a minimum of 4 valid values per group). The mass spectrometry proteomics data have been deposited to the ProteomeXchange Consortium via the PRIDE(78) partner repository with the dataset identifier PXD027029.

### Immunofluorescence analysis

Cells were fixed in 4% paraformaldehyde (PFA, Sigma-Aldrich, Germany) permeabilized with 0.02% saponin (Calbiochem-Merck) and blocked in 0.002% saponin with 2% goat serum (DaKoCytomation, Denmark). The outer membrane of T. gondii was visualized by anti-SAG1 (Abcam, UK) at a concentration of 1/700. As secondary reagents, 1/200 concentrated Alexa Fluor^TM^ 633-conjugated goat anti-mouse IgG (H+L) (Invitrogen, ThermoFisher Scientific, USA) and Cy3-conjugated goat anti-mouse IgG plus IgM (Jackson ImmunoResearch Laboratories, UK) were used. Nuclei were counterstained with 1/2500 4’,6-diamidino-2-phenylindole (DAPI, Invitrogen, ThermoFisher Scientific, USA). The cover slips were fixed in fluorescence mounting medium (Fluoromount-G, Southern Biotechnology Associates, Birmingham, AL). Fluorescence was visualized using a LSM780 confocal microscope (Zeiss, Germany). Image analysis and processing was performed with the ZEN (Zeiss, Switzerland) software.

### Gated Stimulated Emission Depletion measurement

Gated Stimulated Emission Depletion (STED) measurements were performed using a TCS SP8 STED 3X (Leica, Wetzlar, Germany) equipped with an HC PL APO CS2 93x glycerol objective (Leica, Wetzlar, Germany, NA 1.3) at a scan speed of 1400 Hz. For the acquisition of the gated STED signal of the eGFPBoosterAtto488, 488 nm was used as the excitation laser line, 592 nm as the corresponding depletion laser line and the hybrid detector range was set from 498 nm to 580 nm with a 1 ns to 6 ns time gating. For the acquisition of the gated STED signal of the mRFPBoosterAtto647N, 633 nm was used as the excitation laser line, 755 nm as the corresponding pulsed depletion laser line and the hybrid detector range was set from 640 nm to 750 nm with a 0.8 ns to 6 ns time gating. Detection of the different signals was carried out in a frame sequential measurement setup. Deconvolved data of gated STED measurements were calculated using Huygens software (Huygens professional, Scientific Volume Imaging, Netherlands) with reduced signal to noise factor of 5 for eGFPBoosterAtto488 and 6 for mRFPBoosterAtto647N and a reduced iteration setting of 20 for both channels. Resulting deconvolved images were visually compared to the raw data to ensure the quality of the calculation. To correct for channel shift, a reference double staining of nuclear pores was acquired with similar settings and utilized as a reference for channel shift correction by the Fiji-plugin “multistackreg” (79).

### CRISPR/Cas9 knock out strategy

The Galectin 9 gene (lectin, galactose binding, soluble 9, *lgals9*) was disrupted by applying the CRISPR/Cas9 technology in NIH 3T3 fibroblasts (80). The knock out was performed by transfection of the pSpCas9(BB)-2A-GFP (PX458) plasmid (Addgene, USA), expressing a sgRNA (Gal9: for CAC CGC GGG TTA ATG TAT GGA GAC T, rev AAA CAG TCT CCA TAC ATT AAC CCG C) corresponding to the sequence contained in the first exon of the*lgals9* gene. The sgRNA encoding plasmids were transfected into NIH 3T3 fibroblasts according to jetPRIME^R^ transfection protocol (Polyplus, New York City). The transfection efficacy was analyzed by flow cytometry. Subsequently, GFP positive cells were sorted and single cell colonies were cultivated. Mutated clones of the *lgals9* gene were identified by Surveyor® Mutation Detection (Integrated DNA Technologies, Inc., USA) (for AAC TAG ATT GGG CCT GCC TC, rev AGA GAT CCC CCT GAC TCT GT, expected fragments, expected PCR fragment: 923 bp, expected digested fragments: 747, 176 bp). The knockout of *lgals9*was confirmed by PCR amplification, subcloning and sequencing of the corresponding gene stretches from different single cell colonies. Additionally, WB analysis was performed with a polyclonal rabbit anti-mouse antibody against Gal9 (ARP54821_P050, Aviva Systems Biology).

### LDH release assay

Cell death was quantified by measuring LDH release, using the Invitrogen™ CyQUANT™ LDH Cytotoxicity Assay. For this 3x10^4^ cells of the indicated cell lines were stimulated with 100U/ml IFNγ for 16h. Then cells were infected with *T. gondii* ME40 with an MOI of 6.6 for 24h. As positive control, cells were treated with staurosporin for 24h as indicated. % of LDH release was calculated as follows: ((experimental LDH activity- spontaneous LDH activity) x 100) / (maximum LDH activity – spontaneous LDH activity). Each experiment was performed with three technical replicates.

---

**FIGURE S1. Identification and validation of mGBP2 interaction partners.** (A) mGBP2^-/-^ MEFs were reconstituted with HA-mGBP2 or a control vector and stimulated with IFN-γ for 16 h. 1x10^6^ cells were infected for 2 h with *T. gondii* ME49 (MOI 50) or left uninfected. Subsequently, cells were lysed and lysate supernatants were incubated o/n with α- HA antibody coupled agarose beads at 4°C for IP. One part of IP samples was separated via 10% SDS-PAGE and labelled using silver staining. Another part of the IP samples was subjected to Western Botting and immune staining with an α-mGBP2 antiserum or an α-HA antibody. A third part of these IP samples was transferred to the MS analysis (see main text). (B) mGBP2^-/-^ MEFs were reconstituted with GFP-mGBP2 WT as well as one of the N-terminal mCherry fusion proteins mCherry-mGBP2, or mCherry-Gal9. mCherry-Control 1, mCherry Control 2, and mCherry-Control 3 are interacting proteins that will be published elsewhere and do not relate to this study. Cells were stimulated with IFN-γ for 16 h and infected with of *T. gondii* ME49 for 4h. Cells were lysed, and lysate supernatants were incubated o/n with GFP-Trap^®^ beads at 4°C. Pulldown samples and appropriate cell lysate supernatants were subjected to Western Blotting. Blots were stained with α-GFP or α-mCherry antibodies.

**FIGURE S2. Localization of Gal9 and mGBP2 at *T. gondii*.** Recruitment and colocalization of mGBP2 was analyzed in GFP-mGBP2 expressing mGBP2^-/-^ MEFs with additional transduction of either mCherry-Gal9. MEFs were stimulated with IFN-γ for 16 h and subsequently infected with *T. gondii* ME49 for 5 h. After fixation, *T. gondii* were stained with an α-SAG1 antibody and the cell nuclei were labeled with DAPI. Glass slides were analyzed by confocal microscopy. Bars 2 µm.

**FIGURE S3. Colocalization of mCherry-mGBP2 and GFP-mGBP2 at the PV membrane of *T. gondii*.** Recruitment and colocalization of mGBP2 was analyzed in GFP-mGBP2 expressing mGBP2^-/-^ MEFs with additional transduction of mCherry-mGBP2 as control. MEFs were stimulated with IFN-γ for 16 h and subsequently infected with *T. gondii* ME49 for 2 h. After fixation, *T. gondii* were stained with an α-SAG1 antibody. Infected cells were treated with anti-RFP VHH nanobody conjugated to eGFPBoosterAtto647N and with anti-GFP VHH nanobody conjugated to eGFPBoosterAtto488 for enhancement of immunofluorescence of mCherry and GFP respectively. Glass slides were analyzed by STED microscopy. Bars 2 µm. The graphs depict a colocalization analysis of STED images with the ImageJ software (Fiji) for GFP and mCherry fluorescence. The colocalization thresholds were set to 13000-max for the Cherry and 2000-max for GFP in mCherry-mGBP2 and GFP-mGBP2 expressing cells.

**FIGURE S4. Intracellular colocalization of mGBP2 with the interaction partners Gal9**. Colocalization of GFP- mGBP2 with mCherry-Gal9 was analyzed in mGBP2^-/-^ MEFs reconstituted with the indicated fusion proteins. Cells were stimulated with IFN-γ for 16 h. After fixation, cell nuclei were labeled with DAPI. Glass slides were analyzed by confocal microscopy. Bars 5 µm. The right column depicts the results for a colocalization analysis using the Image Visualization and Analysis Software (Imaris) for GFP and mCherry fluorescence.

**FIGURE S5. CRISPR/Cas9 gene editing of Gal9 and verification of Gal9 inactivation.** (A) CRISPR/Cas9 and sgRNA specific for Gal9 were expressed in NIH 3T3 fibroblasts. Clones were picked and DNA from WT and three CRIPR/Cas9 Gal9 targeted cell clones of NIH 3T3 fibroblasts was isolated, PCR amplified (923 bp). PCR products were cloned into the pCR2.1 TA-cloning vector. The Gal9 mutations were verified by Sanger sequencing. Indels, AA sequence and premature stop codons (*) are indicated. (B) For independent mutation analysis, the Surveyor® Mutation Detection Kit for Standard Gel Electrophoresis was employed. Here, a mismatch-specific DNA endonuclease to scan mutations and polymorphisms in heteroduplex DNA is used. PCR amplicons from Gal9 mutant NIH 3T3 clones (test) and WT (reference) DNA were hybridized and the mixtures of hetero- and homo-duplexes were submitted to Surveyor Nuclease digestion. The reference DNA alone, treated similarly, served as a negative control. DNA fragments were analyzed by agarose gel electrophoresis. The formation of new cleavage products, due to the presence of one or more mismatches, is indicated by the presence of additional bands. The relative size of these cleavage products indicates the location of the mismatch or mismatches. (C) Lysate supernatants of NIH 3T3 cells from different CRISPR/Cas9 Gal9 sgRNA targeted clones and from WT cells were analyzed by WB. Cells were stimulated with IFN-γ for 16 h. Blots were stained with α-Gal9 antibodies.

**FIGURE S6. Inactivation of Gal9 does not influence infection rates of *T. gondii***. The infection rates of *T. gondii* ME49 were analyzed in WT and independent NIH 3T3 cell line clones with verified Gal9 inactivation. Cells were stimulated with IFN-γ for 16 h and subsequently infected with *T. gondii* ME49. After fixation, *T. gondii* were stained with the α-SAGI antibody and the cell nuclei were labeled with DAPI. Glass slides were analyzed by confocal microscopy. Bars, 5 µm. The amounts of parasites per cell were quantified 2 h after infection.

**FIGURE S7. Gal9 is required for the intracellular control of replication of *T. gondii*.** WT and independent NIH 3T3 cell line clones with verified CRISPR/Cas9 mediated inactivation of Gal9 (see main text and Figs.S5) were stimulated with IFN-γ for 16 h and subsequently infected with *T. gondii* ME49 for 24 h. After fixation, *T. gondii* were stained with an α-SAGI antibody and the cell nuclei were labeled with DAPI. Glass slides were analyzed by confocal microscopy. Bars, 5 µm.

**FIGURE S8. Cell death of Gal9 deficient cells in *T. gondii* infection.** WT and independent NIH 3T3 cell line clones with verified CRISPR/Cas9 mediated inactivation of Gal9 (see main text and Figs.S5) and NIH 3T3 cell line clones with verified CRISPR/Cas9 mediated inactivation of Gal9 reconstituted with mCherry-Gal9 were stimulated with IFN-γ for 16 h and subsequently infected with *T. gondii* ME49 for 24 h. Subsequently, an LDH release assay was performed. Positive controls were obtained by treating the cells with 1mM Staurosporine for 24h. Shown are three independent experiments, which were respectively performed in technical triplicates to minimize pipetting errors. Plotted are means +/- SD. Statistical analysis was performed using one-way ANOVA followed by Dunnett’s multiple comparison test.; *: p <= 0.05. Only statistically significant comparisons were labelled.

