## Supplementary Figure 1-8 for "GBP2 engages Galectin-9 for immunity against *Toxoplasma gondii*"

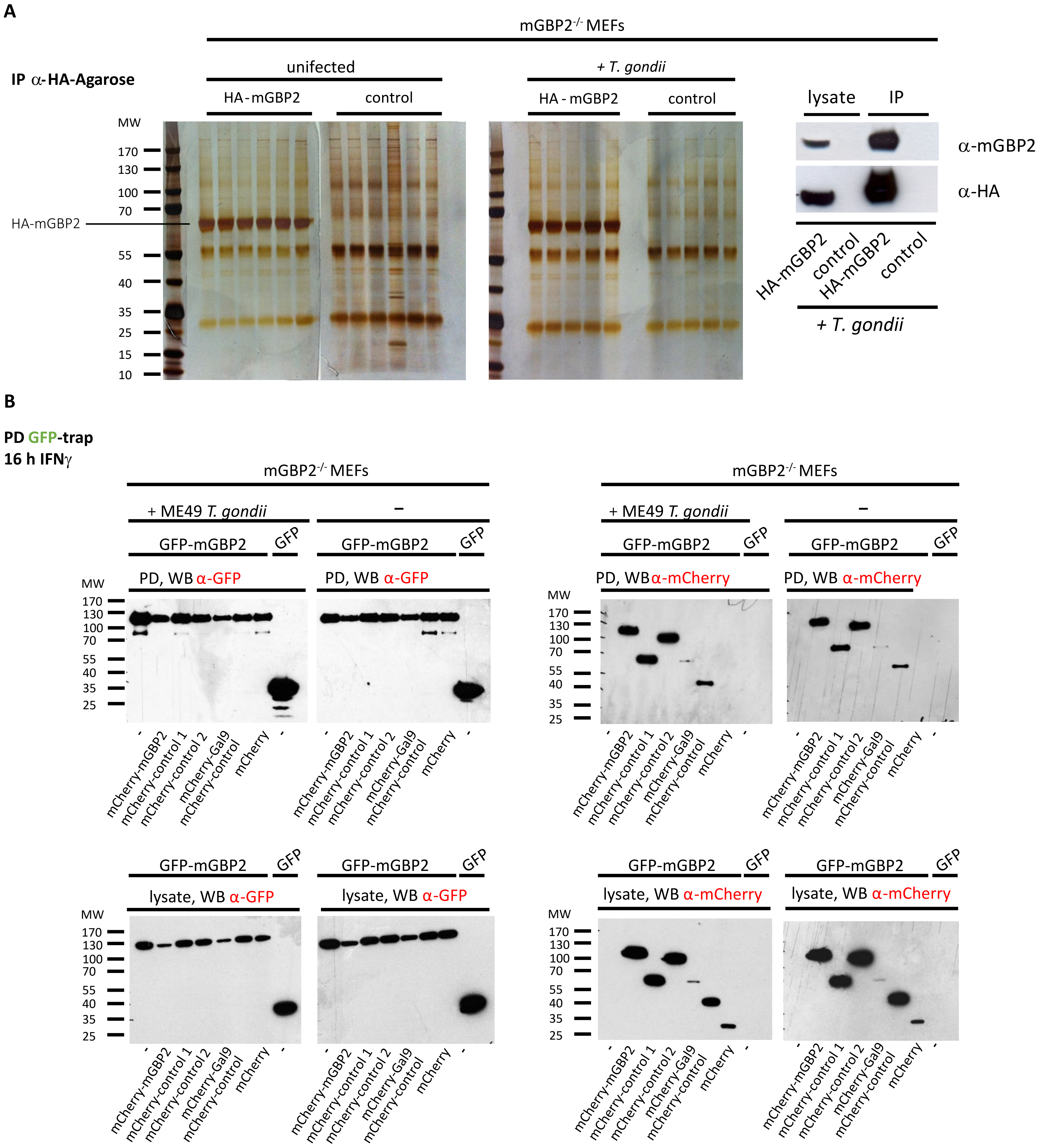
A
IP α-HA-Agarose
mGBP2-/- MEFs
unifected
+ T. gondii
HA-mGBP2
control
HA - mGBP2
control
MW
170
130
100
70
HA-mGBP2
55
40
35
25
15
10
lysate
IP
α-mGBP2
α-HA
HA-mGBP2
control
HA-mGBP2
control
+ T. gondii
B
PD GFP-trap
16 h IFNγ
mGBP2-/- MEFs
+ ME49 T. gondii
–
GFP-mGBP2
GFP
PD, WB α-GFP
PD, WB α-mCherry
lysate, WB α-GFP
lysate, WB α-mCherry
-
mCherry-mGBP2
mCherry-control 1
mCherry-control 2
mCherry-Gal9
mCherry-control
mCherry

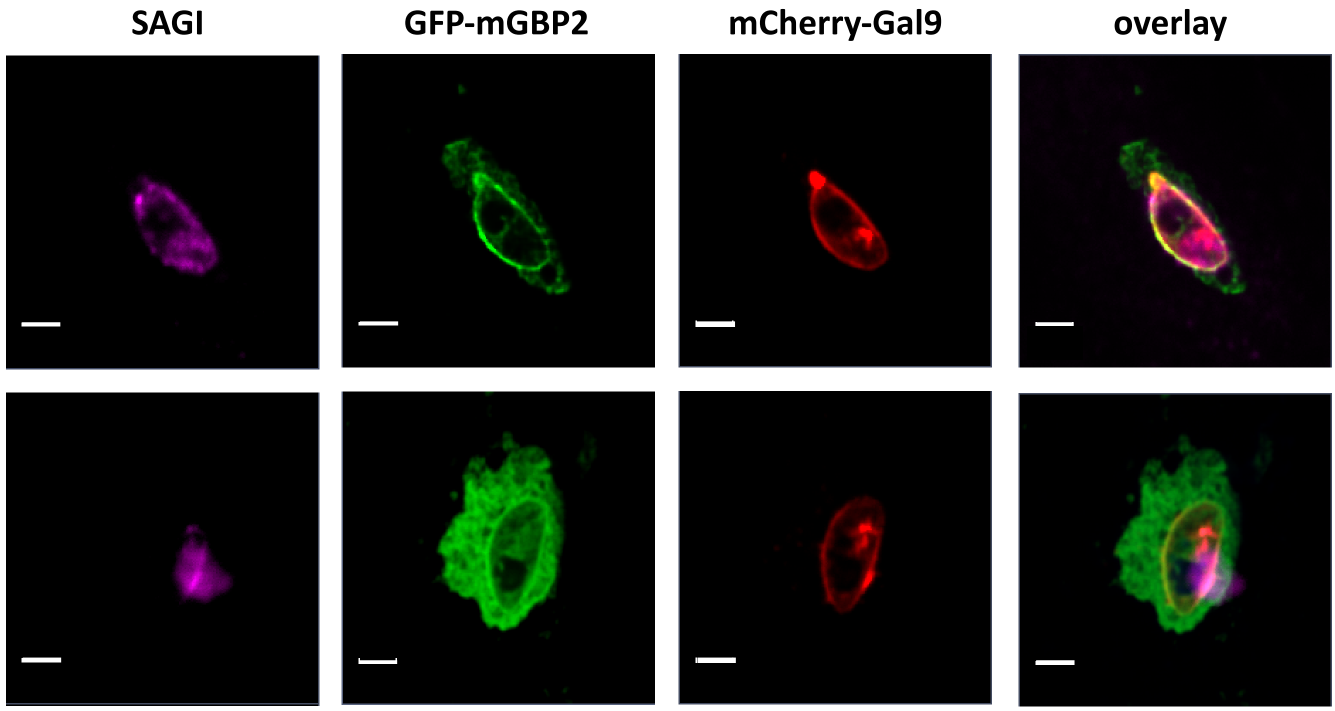
SAGI
GFP-mGBP2
mCherry-Gal9
overlay

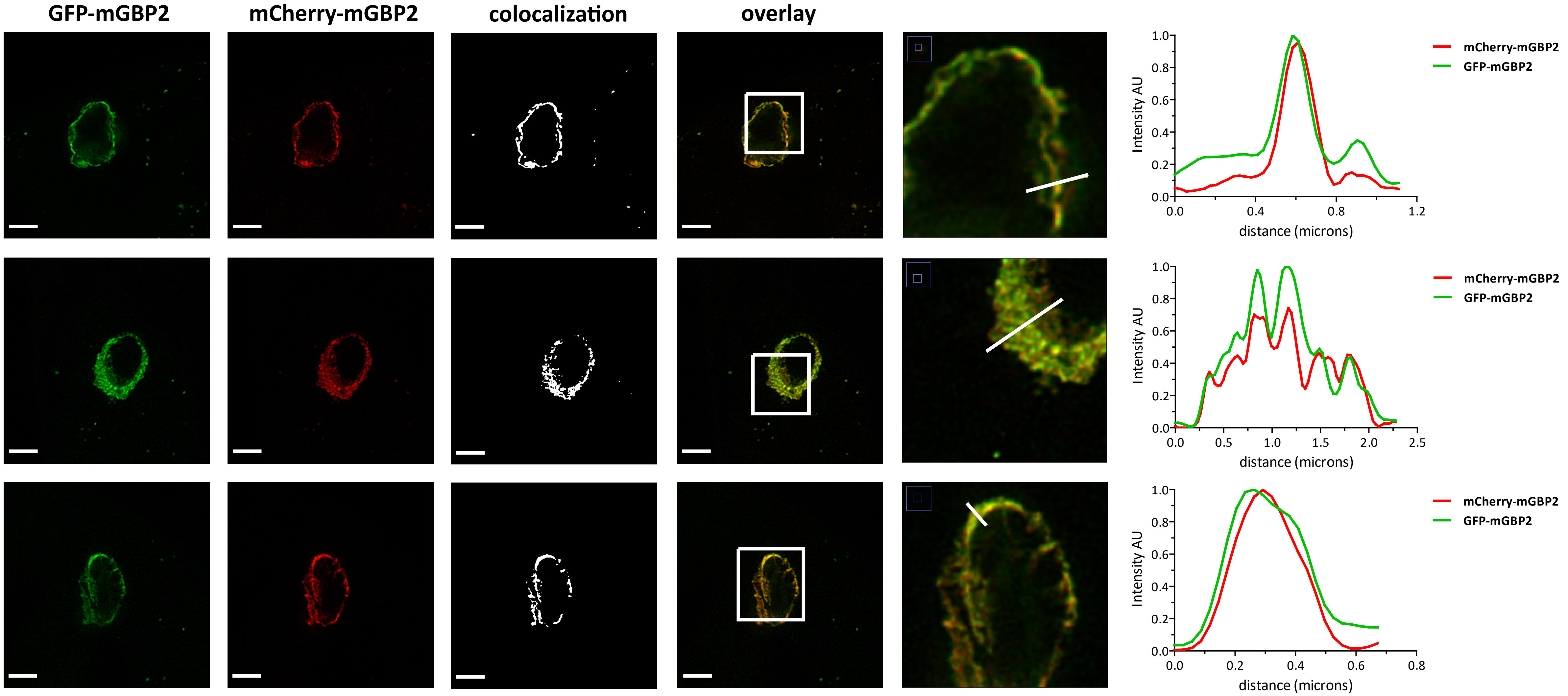
GFP-mGBP2
mCherry-mGBP2
colocalization
overlay
Intensity AU
distance (microns)
mCherry-mGBP2
GFP-mGBP2

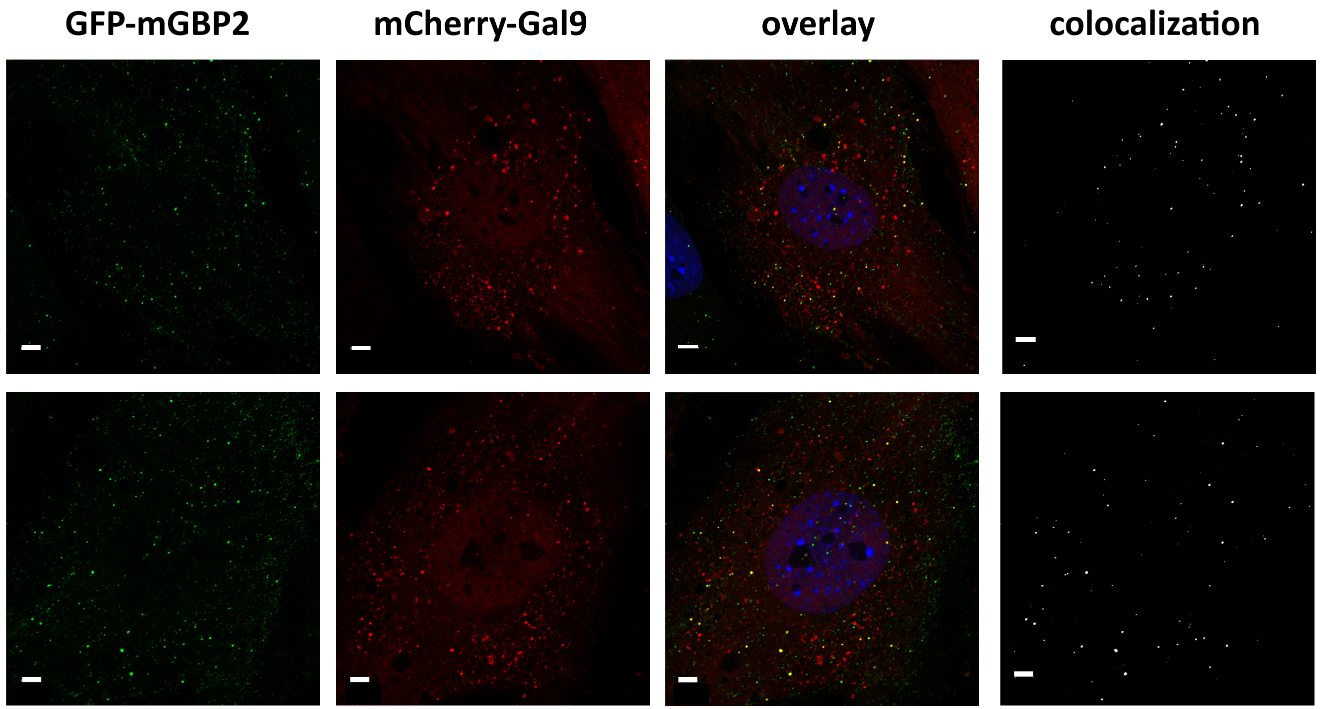
GFP-mGBP2
mCherry-Gal9
overlay
colocalization

A

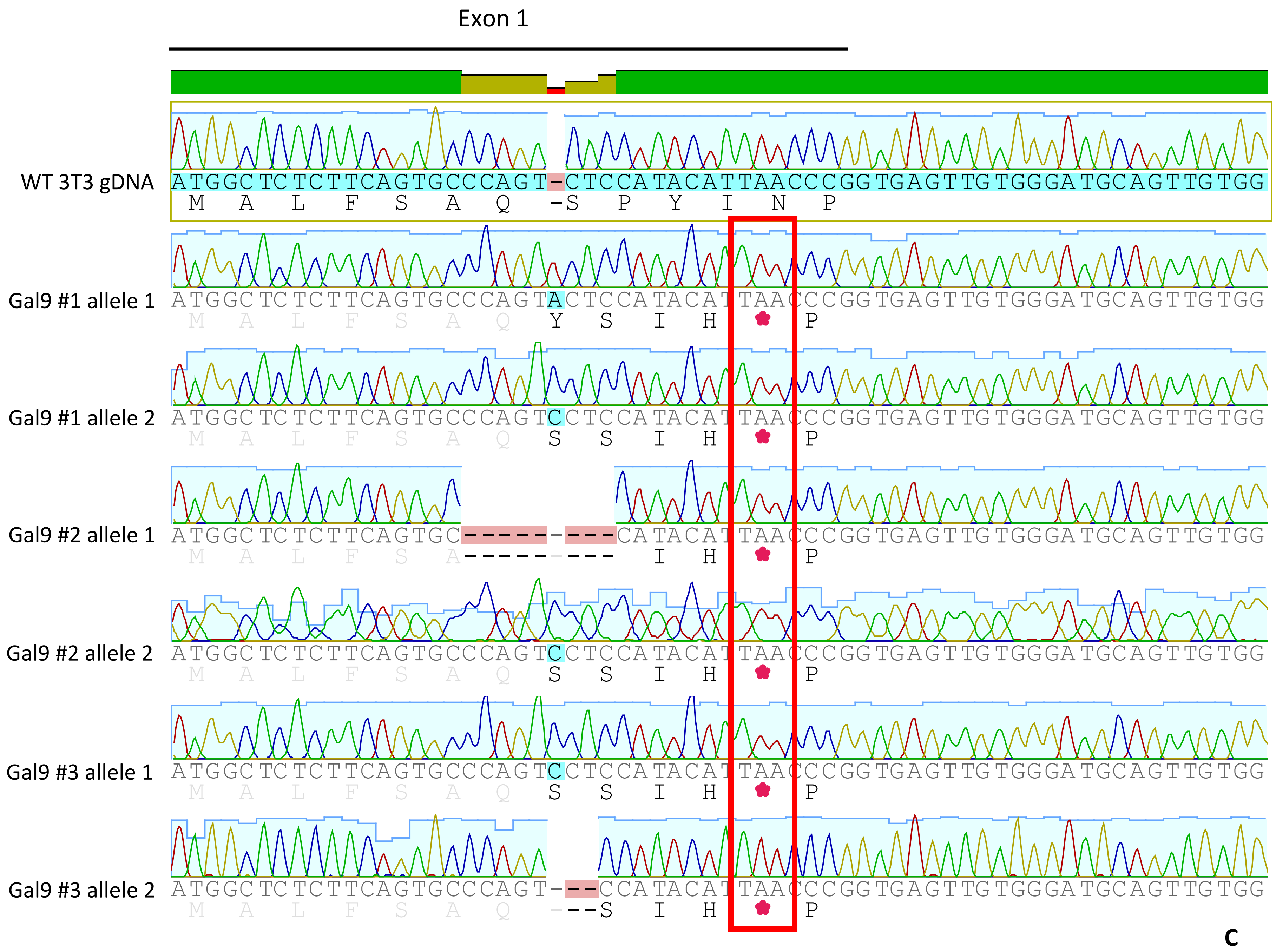


B

| CRISPR/Cas9 | primer fwd | primer rev | PCR fragment (bp) | Surveyor endonuclease fragments (bp) | |
|---|---|---|---|---|---|
| Galectin-9 | AACTAGATTGGGCCTGCCTC | AGAGATCCCCCTGACTCTGT | 923 | 747 | 176 |

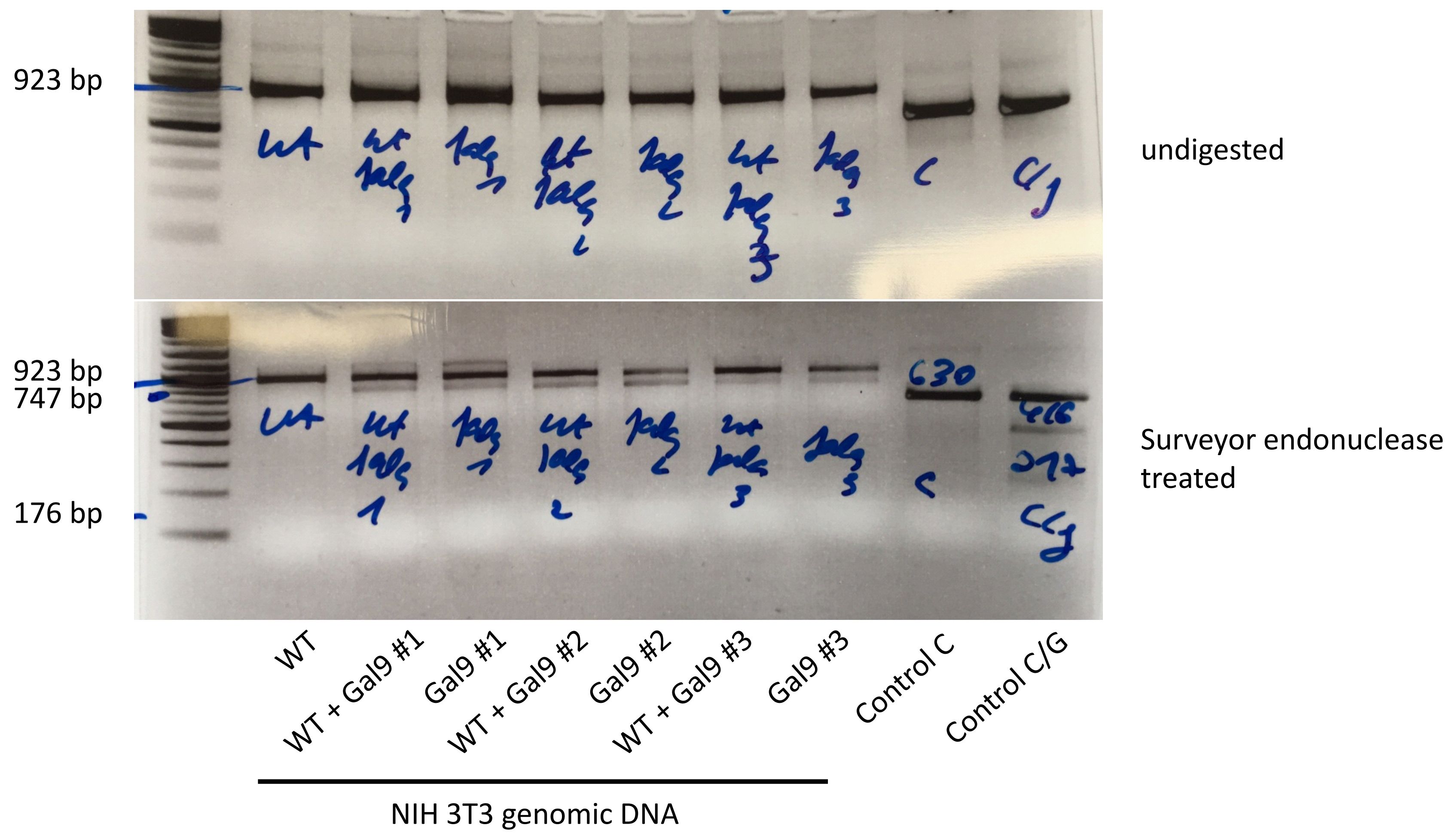


C

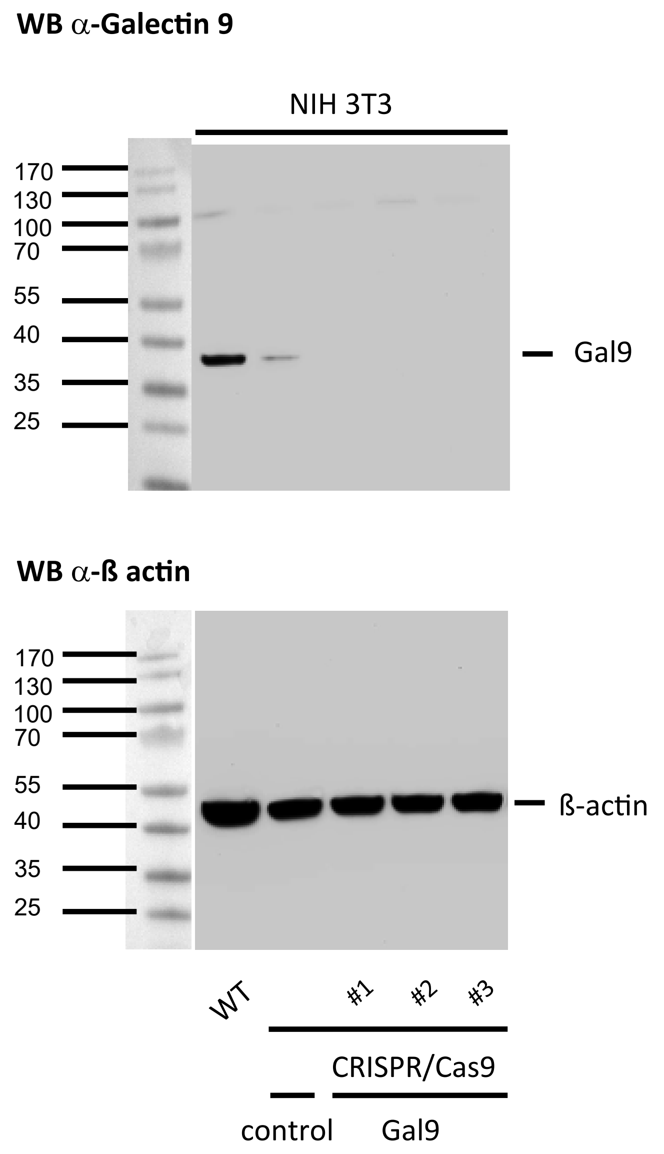

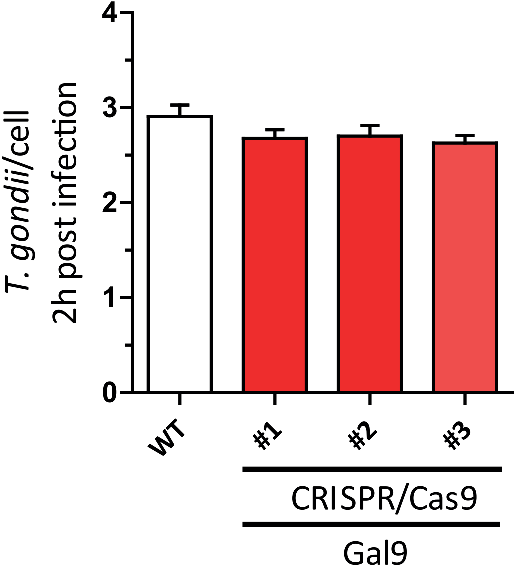
T. gondii/cell
2h post infection
0
1
2
3
4
WT
#1
#2
#3
CRISPR/Cas9
Gal9

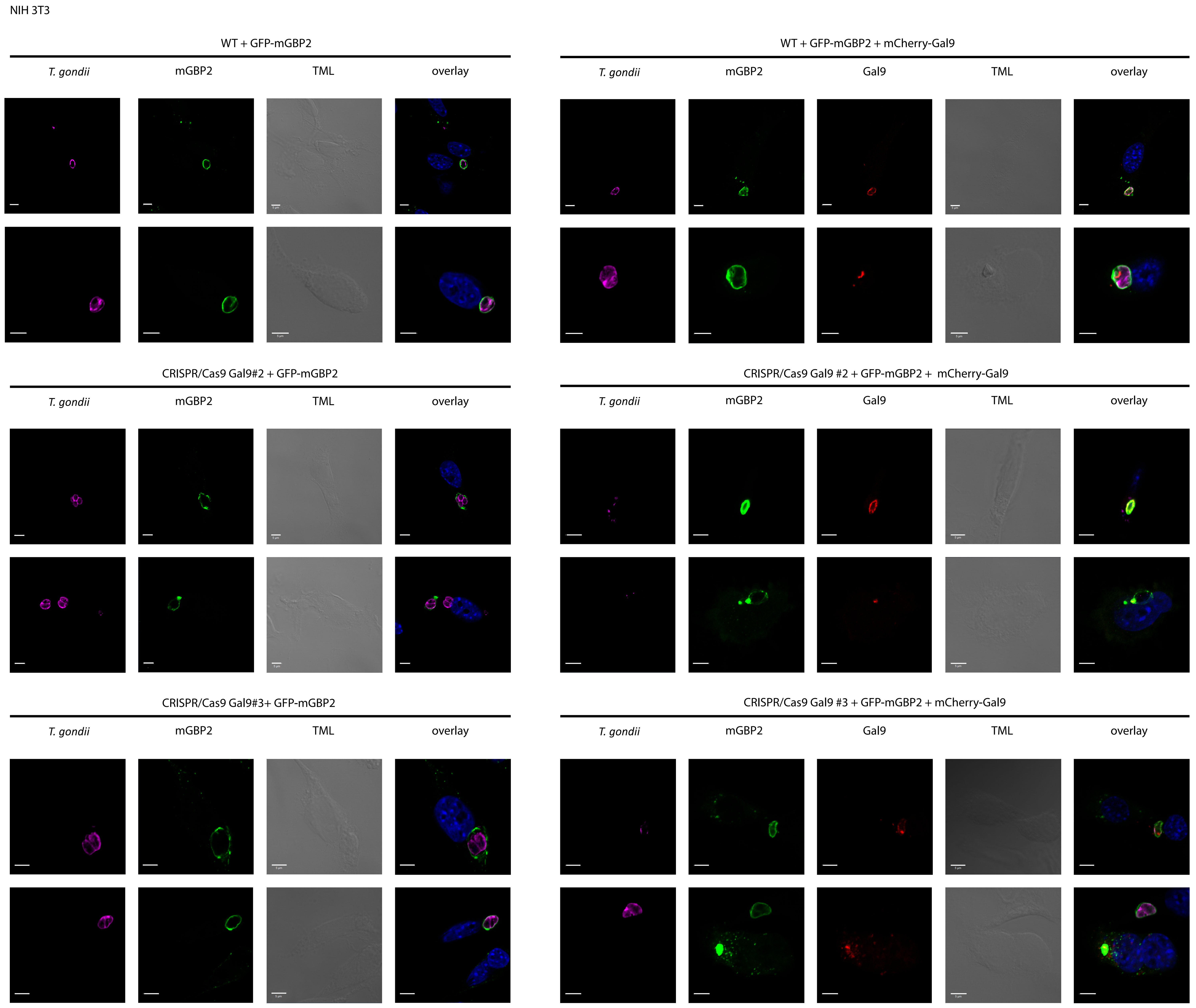
NIH 3T3
WT + GFP-mGBP2
T. gondii
mGBP2
TML
overlay
WT + GFP-mGBP2 + mCherry-Gal9
T. gondii
mGBP2
Gal9
TML
overlay
CRISPR/Cas9 Gal9#2 + GFP-mGBP2
T. gondii
mGBP2
TML
overlay
CRISPR/Cas9 Gal9 #2 + GFP-mGBP2 + mCherry-Gal9
T. gondii
mGBP2
Gal9
TML
overlay
CRISPR/Cas9 Gal9#3+ GFP-mGBP2
T. gondii
mGBP2
TML
overlay
CRISPR/Cas9 Gal9 #3 + GFP-mGBP2 + mCherry-Gal9
T. gondii
mGBP2
Gal9
TML
overlay

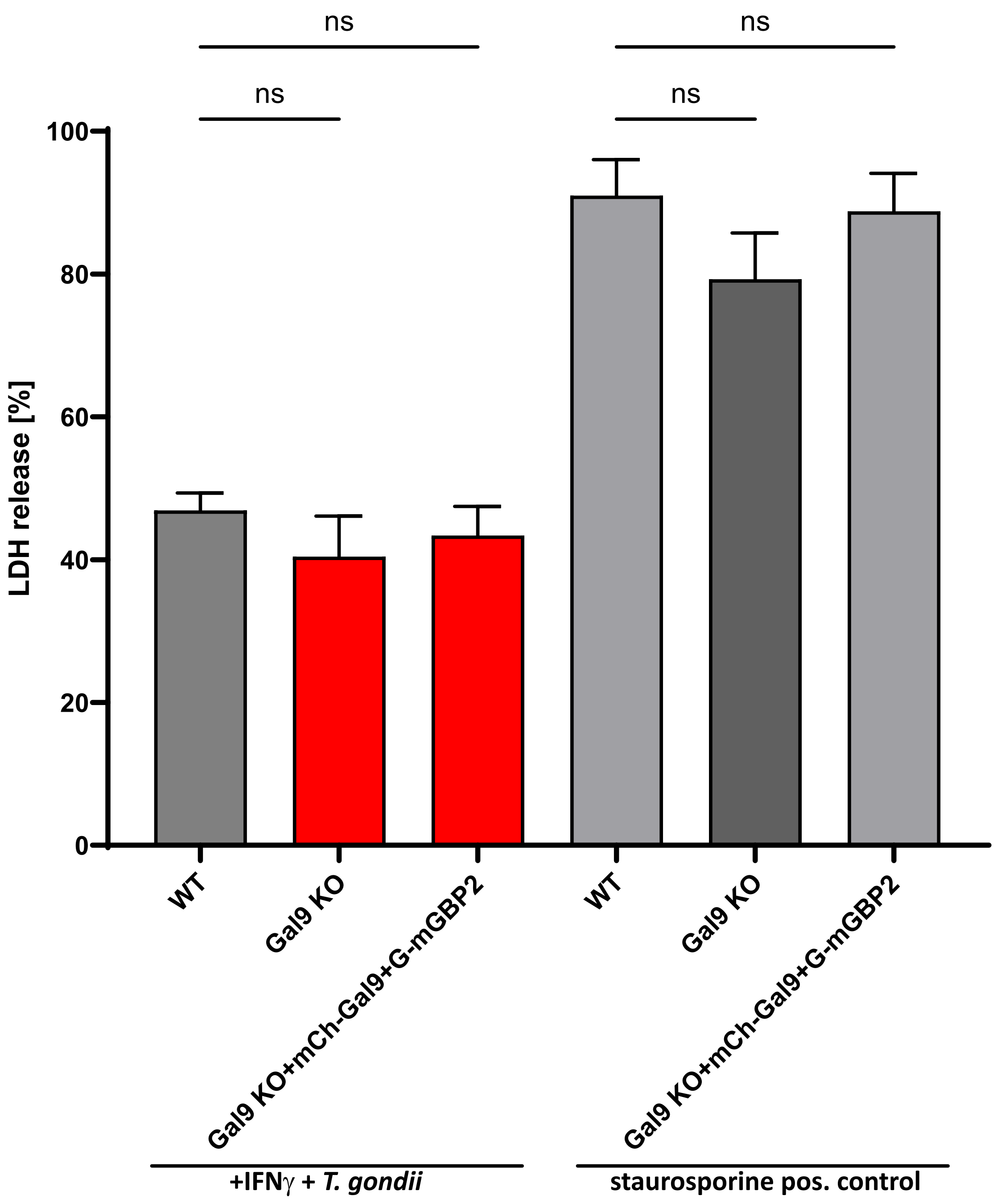
ns
ns
ns
ns
LDH release [%]
100
80
60
40
20
0
WT
Gal9 KO
Gal9 KO+mCh-Gal9+G-mGBP2
WT
Gal9 KO
Gal9 KO+mCh-Gal9+G-mGBP2
+IFNγ + *T. gondii*
staurosporine pos. control
